# On exploring effects of coevolving residues on DNA binding specificity of transcription factors

**DOI:** 10.1101/2021.05.20.445059

**Authors:** Yizhao Luan, Zhi Xie

## Abstract

Transcription factors (TFs) regulate gene expression by specifically binding to DNA targets. Many factors have been revealed to influence TF-DNA binding specificity. Coevolution of residues in proteins occurs due to a common evolutionary history. However, it is unclear how coevolving residues in TFs contribute to DNA binding specificity. Here, we systematically analyzed TF-DNA interactions from high-throughput experiments for seven TF families, including Homeobox, HLH, bZIP_1, Ets, HMG_box, zf-C4 and Zn_clus TFs. Based on TF-DNA interactions, we detected TF subclass determining sites (TSDSs) defining the heterogeneity of DNA binding preference for each TF family. We showed that the TSDSs were more likely to be coevolving with TSDSs than with non-TSDSs, particularly for Homeobox, HLH, Ets, bZIP_1 and HMG_box TF families. Mutation of the highly coevolving residues could significantly reduce the stability of TF-DNA complex. The distant residues from the DNA interface also contributed to TF-DNA binding activity. Overall, our study gave evidence of the functional importance of coevolved residues in refining transcriptional regulation and provided clues to the application of engineered DNA-binding domains and protein.

## Introduction

Transcription factors (TF) regulate gene expression spatiotemporally by binding to regulatory elements of promoter regions of targeting genes and play a central role in genetic activity and human physiology [1, 2]. Decoding DNA binding specificity of TFs is a key to understand the underlying mechanisms of transcriptional regulation of gene expression. Many variables have been revealed to influence the TF-DNA readout on multiple levels, including nucleotide sequence, tertiary structure, TF co-factors, nucleosome occupancy and chromatin accessibility.

Previous studies have provided many important insights into the modes regarding specific DNA recognition by TFs. Studies on TF-DNA structures showed that the preference of a nucleotide at a specific position was largely determined by physical interactions between residues of TFs and base pairs of DNAs, which is also called “base-readout” [3]. For example, ZBTB member ZBTB24 protein interacts with DNA exclusively in the major groove of one 13-bp consensus motif by formation of direct hydrogen bonds and mutation of residues in DNA binding domain would weaken or even lose its DNA binding ability [4]. In addition, TFs could also recognize sequence-dependent DNA structure, such as DNA bending, which is known as “shape-readout” [5]. For example, yeast bHLH TFs Cbf1 and Tye7 bind DNA targets with differential preference for the genomic regions flanking E-Box sites according to DNA shape of binding sites [6]. Another example is Homeodomain TFs Hox-Exd-Hth trimer which prefers DNA sequences with a complex DNA shape that includes optimally spaced minor groove width minima [7]. In the last decade, sequence-based high-throughput technologies to measure protein DNA-binding specificities have revolutionized our ability to measure TF-DNA specificity. The methods based on microarray assays such as protein-binding microarray (PBM) [8, 9], and sequencing assays such as bacterial one-hybrid (B1H) system [10], high-throughput systematic evolution of ligands by exponential enrichment (HT-SELEX) [11] and SELEX-seq [12], enabled large-scale screening of DNA binding preferences of TFs. These studies generally showed that similar TF domains tended to have similar DNA binding sites and different TF members had distinct core binding sites or flanking sequences [9, 13–15]. Nevertheless, many TFs were found to present multiple binding motifs, which makes understanding of the determinants of binding specificity more challenging.

While there is no simple code to specify the interaction of TF and DNA, coevolution of residuals in TFs may help understand factors contributing to TF-DNA binding specificity. The phenomenon that residues in one site change depending on the residues of another site is called “coevolution of residues” [16]. The simultaneous changes in residues have been proven useful for analyzing protein constraints to maintain structural and functional integrity to acquire specific functional necessities [17], understanding protein-protein interaction networks [18, 19], predicting alternative structural conformations and flexibility [20–23] or discovering functional residues that play important roles in catalytic activity and binding affinity of a protein [24, 25]. Regarding coevolving constraints in TF activity, integration of coevolving relationship between TF residues and DNA binding sites could improve prediction of substrate specificity [26–28].

Despite previous studies suggested that coevolving residues may pose structural and functional constraints in several other protein families, whether and to what extent coevolving residues in TF domains contribute to DNA binding specificity is unclear. In this study, we evaluated the effects of coevolving residues in DNA binding domains of TFs on their DNA binding specificity. We systematically collected TF-DNA interactions from high-throughput experiments for nine TF families, including Homeobox, HLH, Ets, HMG_box, Fork_head, bZIP_1, Zn_clus, zf-C4 and zf-C2H2. For each TF family, we defined TF domain sites accounting for TF subclass determining sites (TSDSs). Coevolution analyses revealed that the TSDSs coevolved more frequently with TSDSs than with non-TSDSs. Moreover, we found that some coevolving residues with TSDSs were spatially distant from the DNA interface but impacted stability of TF-DNA complex upon mutation. Our study showed that evolution of residues in TFs played an important role in contributing DNA binding specificity of TFs.

## Materials and methods

### Data collection and processing

We collected TF-DNA interactions obtained with high-throughput technologies (Table S1) for several major species including mouse (Mus musculus), human (Homo sapiens), fruit fly (Drosophila melanogaster), yeast (Saccharomyces cerevisiae) and C. elegans (Caenorhabditis elegans). ChIP-Seq based experiments were not included because of possible confounding raised by TF partners. DNA motifs were presented using position weight matrices (PWMs) [29]. DNA motifs for each TF family were aligned and merged with ‘DNAmotifAlignment’ from motifStack package [30]. TF domain sequences were defined based on the Pfam database [31]. The similarity between DNA motifs was estimated with MotIV package [32]. Multiple sequence alignment (MSA) of TF domain sequences were conducted by MUSCLE (v3.8.31) [33] with the constraint of the typical domain sequence or seed sequence obtained from Pfam database. MSA logos were compared between our collected dataset and the Pfam archives (Figure S1, S2).

### Identification of TF subclass determining sites (TSDSs) of TF

Each TF family was grouped into subclasses by hierarchical clustering analyses based on the pairwise similarity of DNA motifs. Elbow method was used to determine TF clusters. The clusters with within-cluster sum of square lower than 10% of that of the starting cluster without partition were used. The subclasses containing <5 TFs were excluded for the further analysis. Specificity Prediction using amino acid’s Properties, Entropy and Evolution Rate (SPEER) [34], a popular algorithm to identify protein subclass specific sites, was used with the TF clustering relationship as input. In SPEER runs, we assigned equal weights to relative entropy, Euclidean distance, and evolutionary rate of input MSA. The MSA columns with 100% identity in all TF sequences were excluded since they were unlikely to be related to specificity. The residues with p-value <0.1 were defined as TSDSs in our analyses.

### Coevolution analyses of residues

Coevolution analyses of residues were performed using TF domain MSA collected from the Pfam database. Four different algorithms were applied including mutual information (MI) [35], MIp [36], statistical coupling analysis (SCA) [37] and OMES [38]. Specifically, MI measures the reduction of uncertainty in one position by considering the information of the other, thus quantifying between-residues co-variation; MIp is an adjusted MI by removal of background phylogenetic signal of MSA; SCA measures statistical interactions between amino acid positions to map energetic interactions; OMES detects differences between observed versus expected frequencies of residue pairs. All these algorithms were performed using the Evol module of ProDy [39]. Coevolution scores from four algorithms were combined as following: score values between residue pairs from each method were rescaled by formula (*x_i_* - *x_min_*)/ (*x*_max_ - *x_min_*), thereinto, *x*_i_, *x*_min_ and *x*_max_ indicated the score for the *i*-th pair, the minimal and the maximal score respectively; rescaled scores were subjected to quantile normalization; normalized score for each residue pair was then averaged.

Coevolution scores were compared between the groups by Wilcoxon tests, and false discovery rate (FDR) was used to correct the p-values from multiple testing. P value of 0.05 was taken as the cutoff of statistical significance. The coevolving network for each TF family was constructed by taking the residue positions in the MSA as nodes and coevolving relationship as edges. Network analyses and visualization were conducted with igraph (https://igraph.org/). The fast-greedy modularity optimization algorithm was used to detect community structure [40]. To estimate the effects of coevolution on DNA binding specificity, we calculated the similarity scores between the DNA motifs corresponding to TFs containing specific amino acid pair within the CRPs. The DNA motif similarity scores were then summed up using the ratio of specific amino acid pair as weight vector.

### TF-DNA complex structure analyses and computational mutation analyses

TF-DNA structures were firstly collected from the Pfam database. We then downloaded the structures with a resolution <4Å and containing both amino acid and DNA chains from the PDB database (http://www.rcsb.org) [41]. All amino acid chains in PDB structures were included and aligned to the same reference sequence. The distances between protein residues and/or nucleotide were defined using the shortest Euclidean distance between atoms contained in the TF-DNA structure. TF-DNA base contacting and stability of TF-DNA complex upon mutation of amino acid or DNA base were predicted with the protein design tool FoldX [42]. The PDB structures were firstly repaired with the RepairPDB command. Next, phenotypes of DNA mutant were predicted with ‘DNAScan’ command, and that of protein residue mutant were predicted with ‘BuildModel’ command. Foldx simulations were performed for each mutant five-times to increase the conformational space explored, and the averages were reported. Visualization of TF-DNA complex was conducted with Edu PyMol (The PyMOL Molecular Graphics System, Version 1.7.4 Schrödinger, LLC.).

## Results

### Characterization of TF binding specificity

To comprehensively analyze DNA binding specificity of TFs, we collected the publicly available datasets from high-throughput TF-DNA experiments including PBM, B1H or SELEX technology (Materials and methods; Table S1). By a unified data processing strategy, we obtained high-quality DNA motifs for 1,179 TFs from mouse, human, fruit fly, yeast, and *C. elegans*. We discarded the TF families with <30 TF members and finally included DNA binding motifs of 903 TFs in our analysis. These TFs came from nine major TF families where homeobox, zf-C2H2 and HLH were the three largest families (Figure 1A). All these nine TF families were among the top 10 major TF families in animal genomes according to the AnimalTFDB database [43]. Except for Zn_clus family, all the other TF families contained TFs from multiple species. In total, around 85% of TFs were from mouse, human and fruit fly (Figure 1A). In average, around 35.8% of TFs for each family were also found in JASPAR database, which showed high similarity with the annotated ones with a Pearson correlation > 0.9 for each family in average (Figure 1B). Moreover, some TFs showed nearly identical core binding sites comparing with JASPAR collection, such as *NFE2*, *FOXO3*, *USF1*, and *YY1* (Figure 1B). These results suggested the reliability of the datasets.

**Figure 1.**
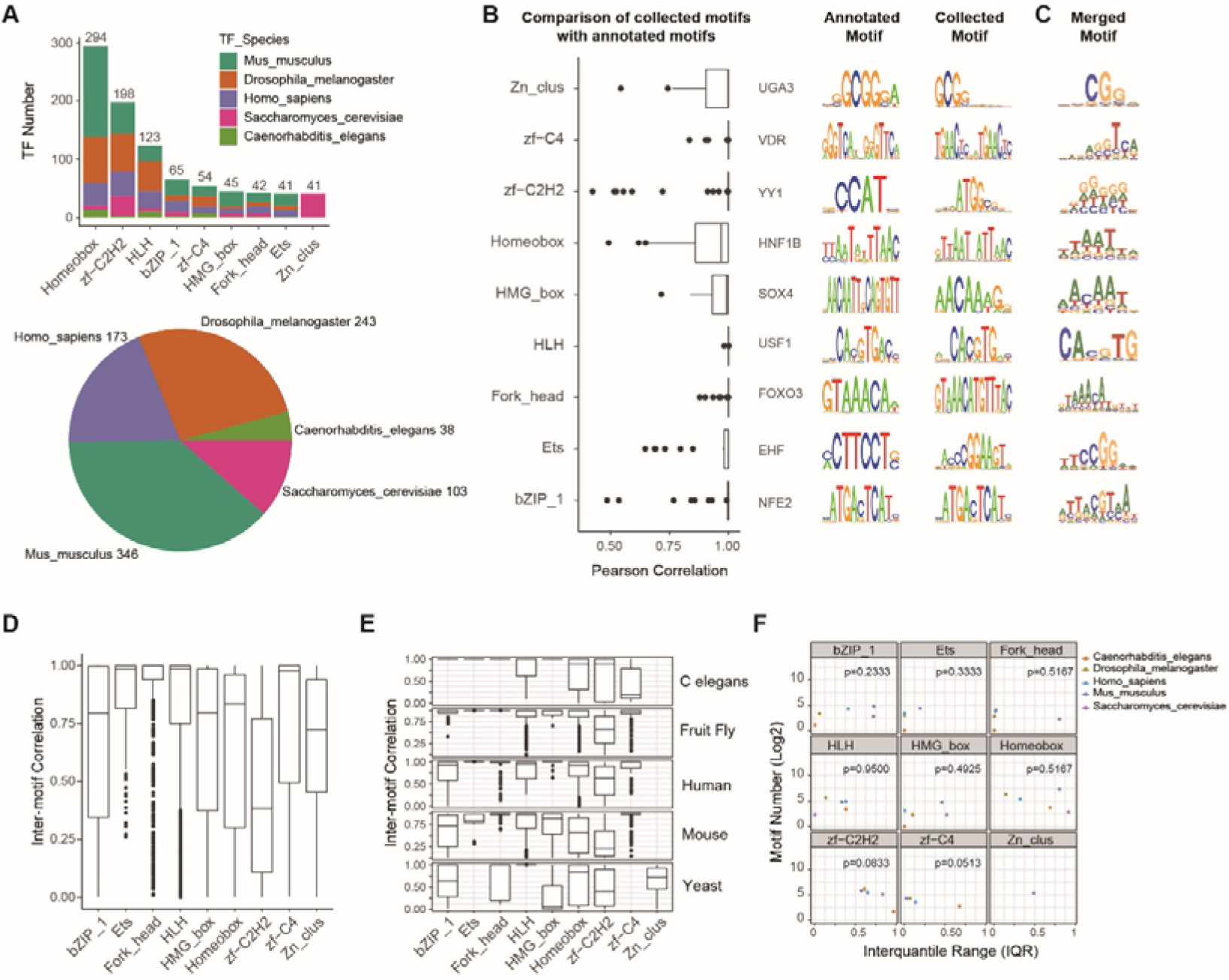
Characterization of TF binding specificity. A. Overview of TF families: (Upper panel) Stacked bar plots showing TF numbers for each TF family where relative percentages of TF in different species are shown with different colors. (Below panel) Pie chart showing total numbers of TF in different species.
B. Boxplot for each TF family (left panel) showing the similarity of overlapped DNA motifs between our included data and JASPAR database. Sequence logos of DNA motifs in JASPAR database (middle panel) and our collected dataset (right panel) of representative TF for each family are compared.
C. Sequence logos of combined DNA motifs in our collected data for each TF family. D, Boxplots showing the similarity of DNA motifs between TFs for each TF family.
D. Boxplots showing the similarity of DNA motifs between TFs in each of five species for each TF family.
E. Scatter plot for each TF family showing no correlation between TF numbers and DNA motif diversity in five species. Spearman correlation analysis and tests were performed.

Aligning DNA motifs for each TF family converged to consensus motifs (Figure 1C). While the degenerated DNA motifs showed distinctive DNA binding sites between TF families, these merged DNA motifs for most families showed changed relative frequency of nucleic acids at core binding sites, suggesting heterogeneity of DNA binding sites in the same TF family. We next calculated inter-motif similarity scores for all pairs of DNA motifs for each TF family. For most TF families including bZIP_1, HMG_box, Homeobox, zf-C2H2, zf-C4 and Zn_clus, the similarity of DNA motifs between TFs varied in a wide range (Figure 1D), indicating common variability in DNA binding sites of TFs even within the same family, consistent with the motif alignment results. Particularly, the similarity of DNA binding sites in zf-C2H2 TFs were significantly lower than the other TF families. Considering TF-DNA interactions we included were from multiple species, the variations of binding preference in a TF family could be caused by difference among species, we further examined the similarity of binding sites within each family for each individual species. A similar pattern was observed (Figure 1E). In addition, we explored whether the variation between DNA motifs was affected by the number of motifs included by calculating spearman correlation scores between inter-quantile range (IQR) of between-motifs similarity and motif numbers for each TF family in each species. For all TF families, no significant associations between higher IQR and motif numbers were observed, which suggested that the higher variation in between-motifs correlation was unlikely caused by greater number of motifs (Figure 1F). There results together demonstrated that DNA binding preference was divergent between different TF families and even TFs within the same TF family exhibited different degrees of heterogeneity in DNA binding specificity.

### Identification of TF subclass determining sites (TSDSs)

We next sought to determine the TSDSs of TFs based on their DNA binding site patterns. For each TF family, we firstly divided TF members into different subclasses based on the similarity of DNA motifs (Materials and Methods). We noticed that the zf-C2H2 contained more than 20 subclasses which made the number of TF members in the subclasses too small, which was consistent with the observation in another recent systematic analysis on TFs [44]. This is because that zf-C2H2 TFs usually contain multiple copies of DNA binding domain, which bind DNA as an array which allows these TFs to recognize new binding sites [1, 45]. On the other hand, the Forkhead TFs had high similarity between DNA motifs, where only one subclass was identified. We therefore discarded these two TF families from further analysis.

The remaining seven TF families were sub-grouped based on the diverged core or flanking sequences of binding targets. Of note, we identified TSDSs or major TF subclasses consistent with previous studies on Homeobox, HLH, Ets, HMG_box, and bZIP_1 TFs (Figure 2). The TSDSs for zf-C4 and Zn_clus TFs are shown in Figure S3. 62% Homeobox TFs bind typical DNA motif ‘TAAT’ [46]. The residue positions 53, 46, 49, 54 and 25 in the Homeobox domain were identified to be associated with different DNA motifs. Among these five sites, four positions (53, 46, 49, and 54) were on the recognition helix. Moreover, the residue 49 had been experimentally proved to be crucial in TF-DNA binding specificity by mutation assays [47]. For the HLH family, almost all the included TFs preferred to recognize DNA sequence ‘CANNTG’, known as “E-box” [48]; but four positions (5, 8, 13 and 14) on the HLH domain correlated with different forms of E-box, consistent with previous studies. For example, Arg13R was used to specify CACGTG motif, in line with the inference by half-sites-based analysis [49]. For Ets TFs, our analysis revealed that five amino acid positions (31, 51, 53, 61, and 76) correlated with DNA binding specificity where the positions 51 and 53 were on helix-3 of the domain and the position 76 was on strand-4 [50]. For HMG_box TFs, five amino acid positions (19, 20, 23, 27, and 49) were found to be associated with DNA motifs clusters where the positions 19, 20, and 23 were on alpha helix 1, and the position 27 was at the N terminus of alpha helix 2, known as typical structures of HMG-box domain [51]. For bZIP_1 TFs, seven amino acid positions (4, 8, 14, 17, 18, 20, and 31) correlated with DNA motif clusters where the positions 17 and 20 were reported to be signature for DNA recognition [52]. These results together indicated that our analysis revealed reliable relationship between TF subclass and their specific DNA binding activity.

**Figure 2.**
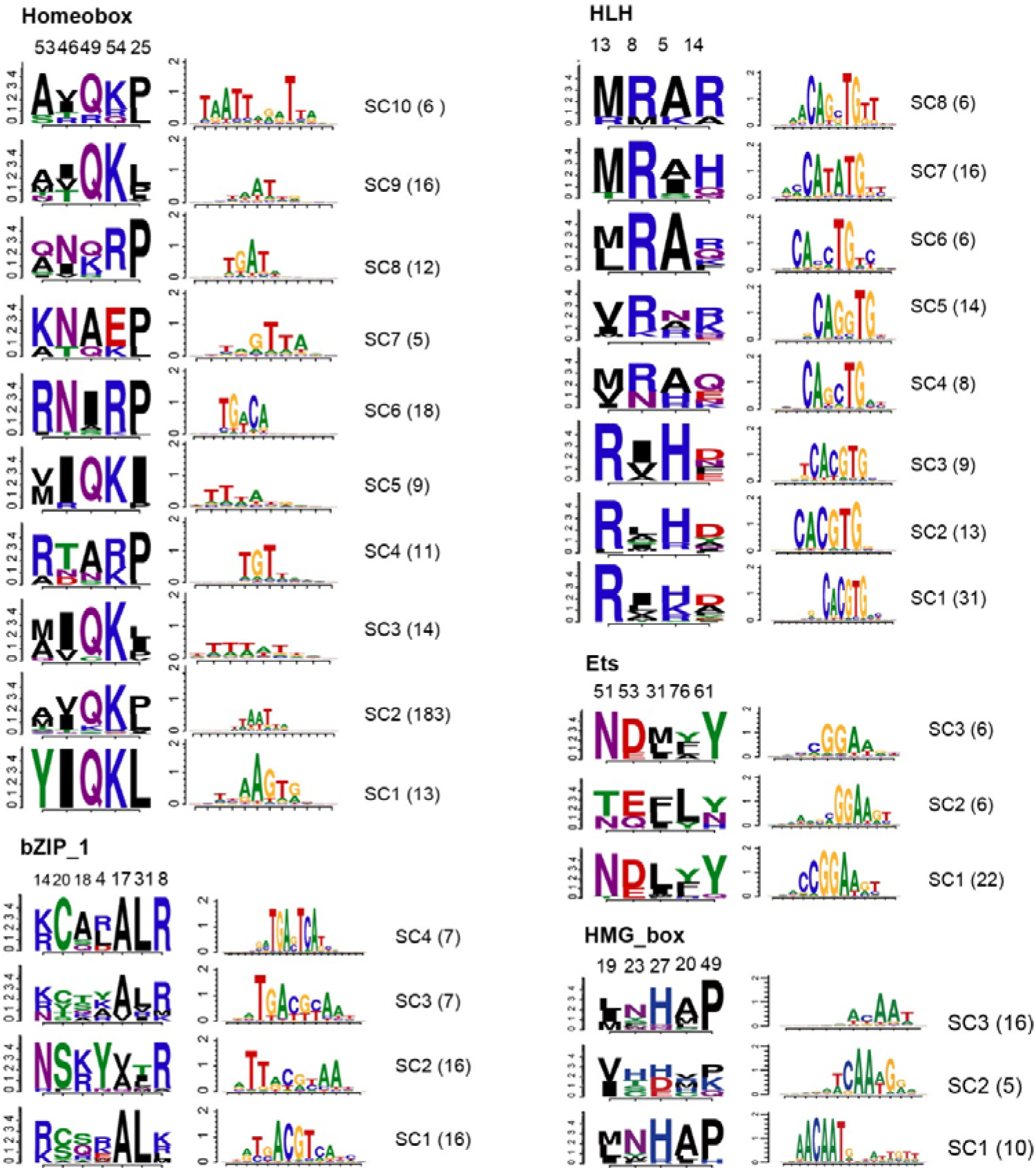
Identification of TF subclass determining sites (TSDSs). Sequence logos of TSDSs (left panel) and corresponding merged DNA motifs (right panel) for each TF subclass from five TF families: Homeobox, HLH, bZIP-1, Ets and HMG_box. The number of members of each subgroup is shown in the parenthesis.

We further found that DNA specificity usually correlated with more than one TSDS positions. For example, our results revealed that DNA specificity of Homeobox TFs was not limited to the well-studied amino acid position 49, the residues 53R and 49Q were observed in different TF subclasses corresponding to DNA motif clusters. Take HLH TFs as another example, 13R and 8R also appeared to be mutually exclusive, and combinations of 8R and the amino acid at the other positions accounted for the divergence of non-canonical E-box, such as CATGTG and CAGGTG. These results suggested that TSDS combination could predict DNA binding targets more accurately, and TSDS positions tended to covary in terms of amino acid composition.

### Coevolving residue pairs (CRPs) and TSDSs

Correlated mutations or covariation between residues were thought to be suggestive of coevolution [53], we next explored whether and to what degree these TSDSs coevolved corresponding to DNA binding specificity. To this end, we collected protein sequence alignments of various species from the Pfam database. These protein sequences were re-aligned to the same seed sequence for each TF family. We developed an ensemble strategy by applying multiple methods including MI, MIp, SCA and OMES to detect reliable CRPs (Materials and Methods). For each method, CRP candidates were defined as the top 10% residue pairs according to the estimated measurements. In general, MI, MIp and OMES yielded moderately to highly consistent results with Jaccard similarity coefficients ranging between 0.43~0.83, while SCA measurement weakly correlated with all the other methods (Figure S4), consistent with a benchmarking study on coevolution methods [54].

CRPs were further defined with the candidates identified by at least two methods for each TF family (Table S2). We observed CRPs between TSDSs and/or non-TSDSs. As non-TSDSs accounted for most of the residues in TF domains, around 81% CRPs were among non-TSDSs in average while around 2.7% CRPs were among TSDSs in Homeobox, HLH, bZIP_1, Ets, and HMG_box TFs (Figure 3A). Interestingly, in these five TF families, we found that TSDSs tended to be more frequently coevolving with TSDSs, revealed by network-based community analysis showing that our identified TSDSs were usually clustered together (Figure 3B; S5). To understand the influence on the DNA binding specificity of CRPs, we compared the similarity of DNA motifs of TFs grouped by CRPs or by non-CRPs (Materials and Methods). Interestingly, we found that TFs grouped by CRPs had higher degree of DNA motif similarity than that by non-CRPs in the six TF families out of seven we tested, except for Ets TFs (Figure 3C), suggesting that the CRPs were related to similar DNA binding activities.

**Figure 3.**
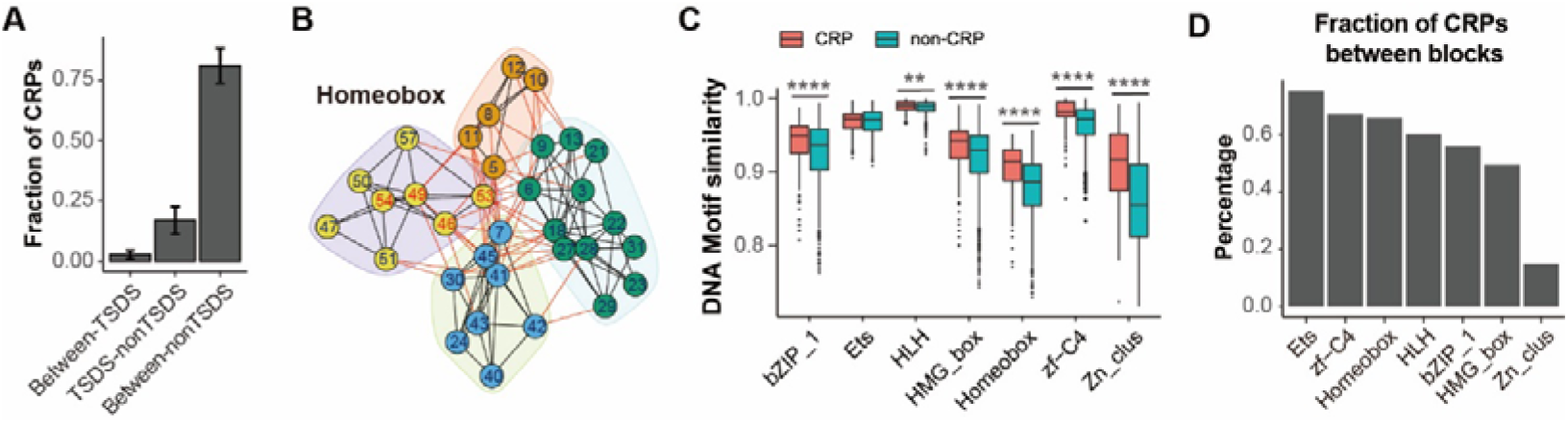
CRPs and TSDSs. A. Bar plots showing the fraction of CRPs between TSDSs, between TSDSs and non-TSDSs and between non-TSDSs across all TF families.
B. Representative Network-based partition of coevolving residues in Homeobox. Nodes and numbers refer to residue index in TF domain, edges refer to coevolving relationship. Numbers in red indicate TSDSs. Nodes in different clusters are shown in different colored background.
C. Boxplots showing comparison of DNA motif similarity of TFs grouped by CRPs and non-CRPs. (**), p<0.01; (****), p<0.0001.
D. Bar plots showing the fraction of out-of-block coevolution for each TF family.

We next examined whether the CRPs were adjacent in the sequence of amino acids, known as blocks that were considered to be important for protein evolution [55]. We found that more than half of the CRPs were between residues more than five positions apart from each other in the alignment of six TF families, except for the Zn_clus TFs (Figure S6). By defining the residue blocks (Table S3), we further found that as high as 75% CRPs were between different blocks in bZIP_1, Ets, HLH, HMG_box, Homeobox, and zf-C4 TFs, while the out-of-block coevolution only accounted for ~15% CRPs in Zn_clus TFs (Figure 3D), suggesting that the CRPs were not always continuous in TF domain sequence positions.

### CRPs and TSDSs in TF-DNA complex

In addition to residue blocks conveying coevolution constraints, it is known that TF residues located in the TF-DNA interface are also likely to impact DNA binding [45]. We therefore sought to understand how these CRPs and TSDSs locate in the TF-DNA complex. We totally collected 178 available TF-DNA complexes (Table S4) from the PDB database for seven TF families, of which the side chains containing DNA binding domains were aligned individually. Mapping TF domain residues to these 3D structures, we estimated the spatial distance of all pairs of residues in the TF domain (Materials and methods). As expected, the CRPs located closer in 3D structures than the non-CRPs for all the TF families (T-test, p<10^−8^ for all TF families; Figure S7). Nevertheless, we noticed that around 31% of CRPs in average across all seven families had a spatial distance >10□, consistent with previous findings that not all the coevolving residues were close in the 3D structures [56]. We next analyzed whether each residue in CRPs was close to the TF-DNA interface that could present a constraint on their coevolution. Comparing the distance to DNA interface of each residue in a pair of coevolving residues, we found that in all seven TF families, the percentages of CRPs that had at least one residue far away from the DNA interface (f-CRPs), with a distance >10□, ranged from 18.6% to 67.5%, with a median of 33.3% (Figure 4A), suggesting a heterogeneous relationship between coevolving residues of TF domain and DNA binding activity in different TF families. Among these f-CRPs, most were between non-TSDSs while several were between TSDSs and non-TSDSs in Homeobox, HLH, Ets, bZIP_1 and HMG_box TFs. Of note, the distance to DNA of the TF residue off the interface varied in a wide range, as far as > 50□ in bZIP_1 TFs (Figure 4A).

**Figure 4.**
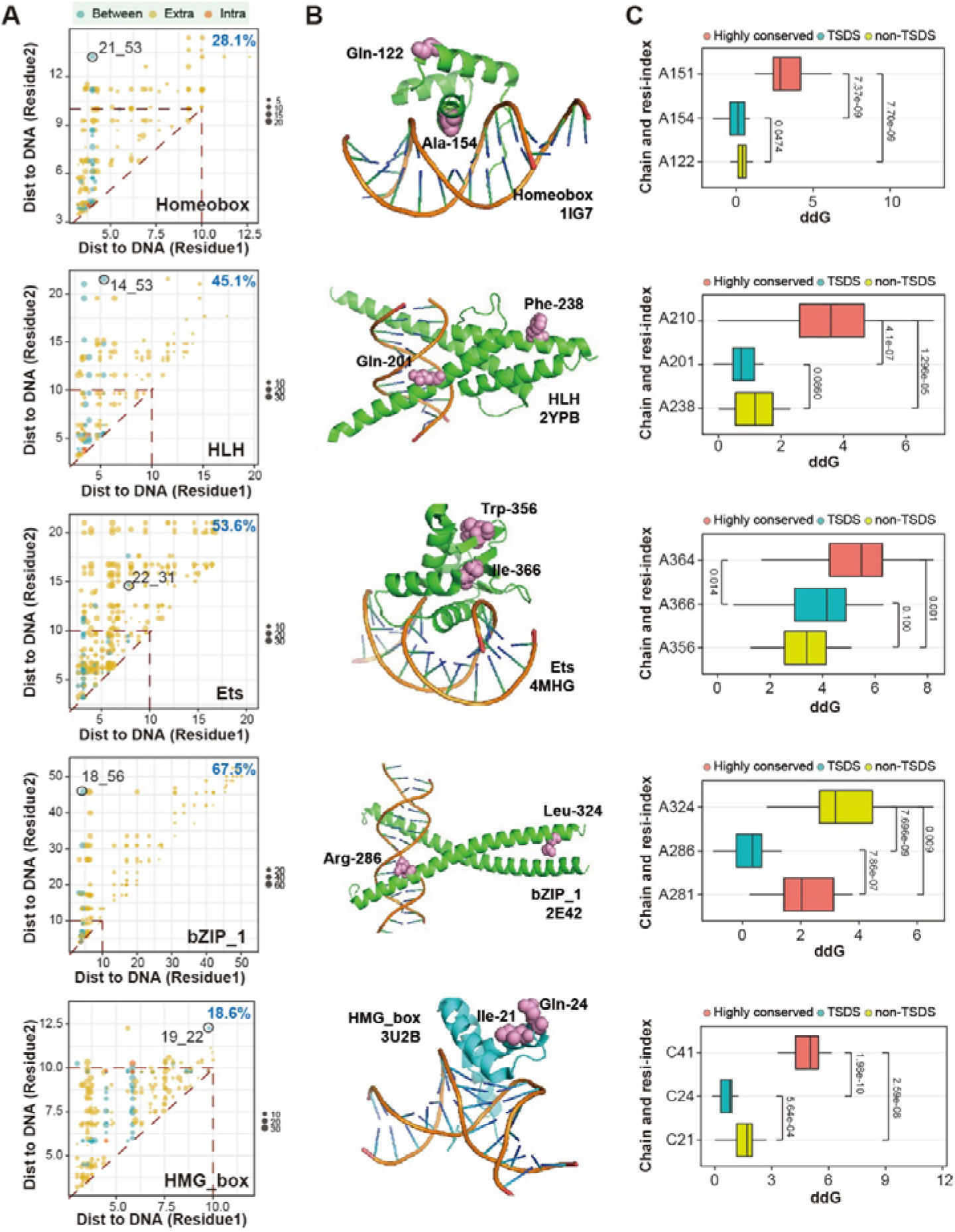
CRPs and TSDSs in TF-DNA complex. A. Scatter plots showing comparison of distance to DNA interface of each residue in CRPs in Homeobox, HLH, Ets, bZIP_1 and HMG_box TFs. The CRPs from different group of residue pairs between-TSDSs (intra), between TSDSs and non-TSDSs (between) and between-non-TSDSs (extra) are in different colors. For each TF family, the percentage of CRPs having at least one residue with a distance>10A to DNA are calculated. Representative CRPs between TSDSs and non-TSDSs are highlighted in circles and noted with residue index.
B. Representative PDB structures for Homeobox (1IG7), HLH (2YPB), Ets (4MHG), bZIP_1 (2E42) and HMG_box (3U2B) families. Representative CRPs shown in panel A are highlighted in red spheres and noted with amino acid type and residue ID within one side chain containing TF domain.
C. Boxplots showing the ΔΔG of mutants of indicated residues in selected CRPs by comparing with wild type of selected PDB structures in Homeobox, HLH, Ets, bZIP_1 and HMG_box TFs. Side chain of amino acid and residue IDs are used to indicate the residue mutant. One-tail Wilcoxon tests were conducted in statistical testing.

To analyze how the residues in f-CRPs impacted DNA binding, we estimated the changes of interaction energy (ΔΔ*G*) upon mutation of these residues in TF-DNA structures (Materials and Methods). We conducted *in-silico* modeling with mutation analyses on selected f-CRPs from representative PDB structures (PDB: 1IG7 for Homeobox, PDB: 2YPB for HLH, PDB: 4MHG for Ets, PDB: 2E42 for bZIP_1 and PDB: 3U2B for HMG_box) from the five families in which we identified known TSDSs (Figure 4B). Each CRP was between TSDS and non-TSDS (Figure 4A), thus we could compare the impact of mutations of TSDS and non-TSDS on TF-DNA binding activity. We also conducted mutation analysis of highly conserved amino acid sites in these TF families and compared with that of CRPs. We found that mutations of highly conserved residues induced significantly higher ΔΔ*G* than the other tested mutations in five out of six TF families including Homeobox, HLH, Ets and HMG_box (Figure 4C). Of note, we found that mutant of non-TSDS residue off the DNA interface induced significantly higher ΔΔ*G* than that of TSDS residue close the DNA interface in Homeobox, HLH, bZIP_1 and HMG_box TFs (Wilcoxon test, all p<0.1). In addition, in Ets TFs, mutation of either TSDS or non-TSDS induced a ΔΔG>2 kcal/mol, which was generally thought to be enough to completely disrupt the DNA binding capabilities [49]. These results together demonstrated the biological effects on DNA binding of coevolving TF residues, even for those spatially distant residues from the DNA interface.

## Discussion

In this study, we conducted a comprehensive and integrative analysis of coevolution of residues in TFs and their DNA binding motifs for seven TF families. We provided evidence showing that coevolving residues in TF domains contributed to DNA binding specificity. We demonstrated that the TSDSs were more likely to be coevolved with TSDSs than with non-TSDSs. Mutation of the CRPs could significantly reduce the stability of TF-DNA complex and even the distant residues from the DNA interface contribute to TF-DNA binding activity.

Our analysis identified multiple TSDSs consistent with the known determinants in five out of seven TF families, including Homeobox, HLH, Ets, bZIP_1 and HMG_box TFs. We recovered the specific patterns of DNA binding site for several known TSDSs. We also noticed that our TSDSs did not recover all the known amino acid sites related to DNA binding specificity, such as the flexible N-terminal arm of homeodomain which can show base-specific contacts with the minor groove via conserved arginine in this region [57]. The reasons could be that our analyses were based on monomers of DNA binding domains and our analyses excluded highly conserved residues from the domain MSA profiles. Further studies will be conducted by integrating complex interaction between TF domains such as homodimers and heterodimers of TFs.

We showed the importance of coevolving residues in structural integrity and DNA binding specificity of TFs with *in-silico* mutation analysis on representative TF-DNA structures from five TF families. Notably, our analysis revealed the residues distant from the DNA interface but showing considerable impacts on DNA binding compatibility upon mutation. For example, the residue LEU324 in CEBPB protein is distant from the bound DNA but can make the DNA binding complex incompatible upon mutation to the other amino acid types (Figure 4). This may be part of the reason to those coevolving residues in TF domains but not contacting with each other.

To sum up, our studies expand our knowledge on the interaction between coevolved residues in TFs, tertiary contacting, and functional importance in refined transcriptional regulation. Understanding the impact of coevolving residues in TFs will help understand the details of transcription of gene regulation and advance the application of engineered DNA-binding domains and protein.

## Acknowledgements

This work was supported by the National Natural Science Foundation of China (31900477 to YZ.L.), and China Postdoctoral Science Foundation (2018M633217 to YZ.L.).

## Authors’ contributions

Z.X. conceived and designed the project. YZ.L. and Z.X. analyzed the data and wrote the manuscript. All the authors read and approved the paper.

---

**Figure S1.**
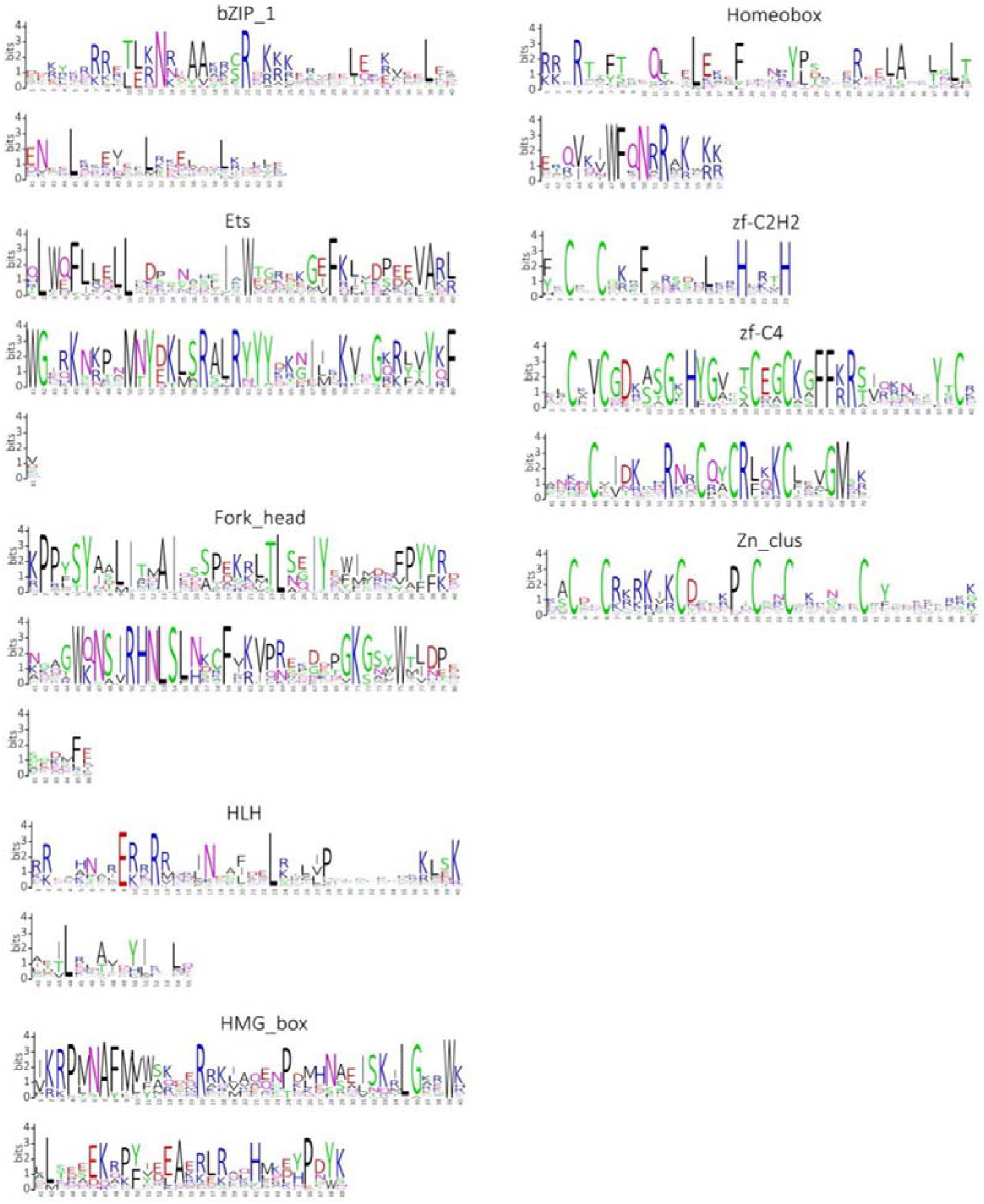
Sequence Logos of aligned DNA binding domains in our analysis for each TF family.

**Figure S2.**
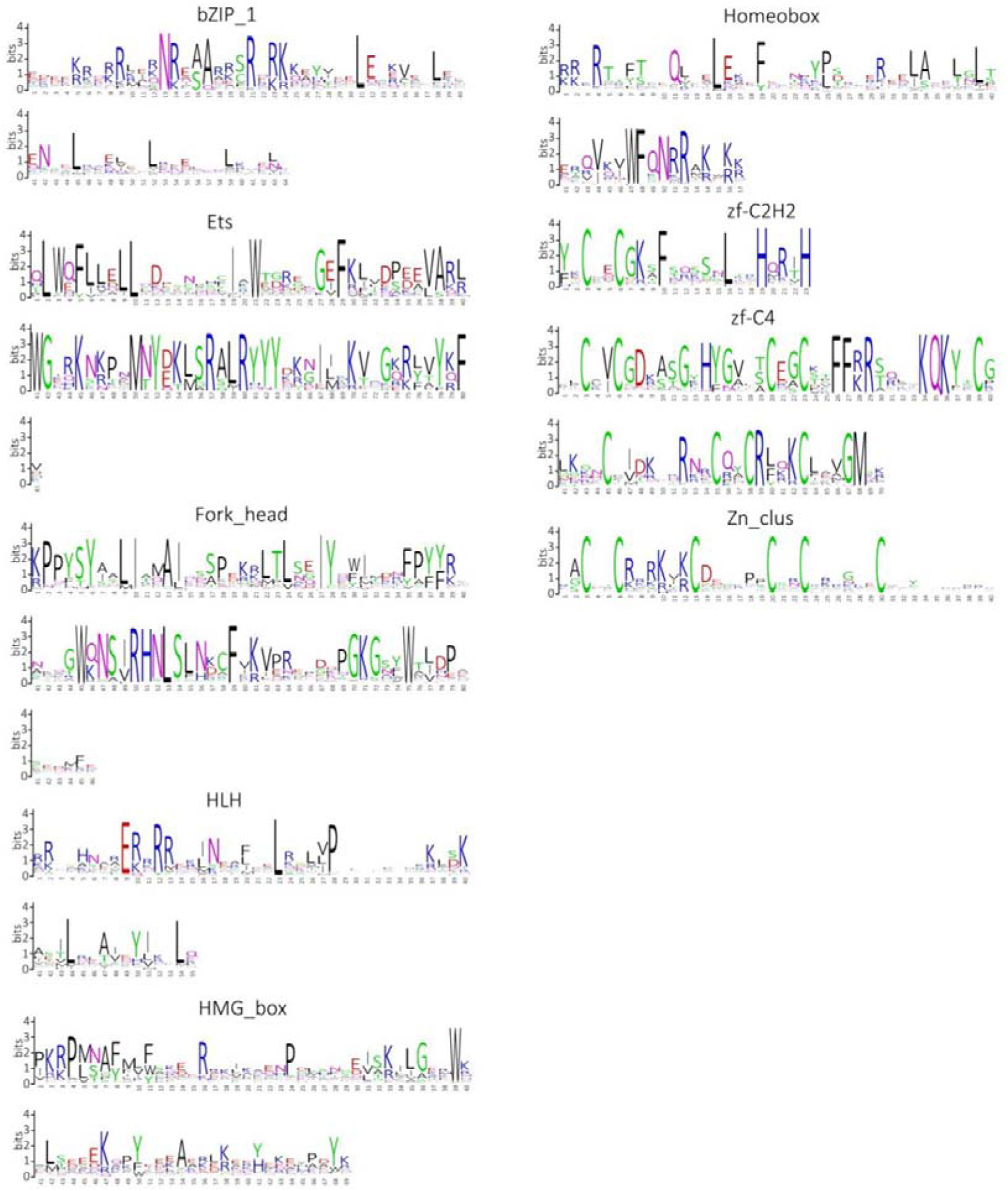
Sequence Logos of DNA binding domains for each TF family in Pfam.

**Figure S3.**
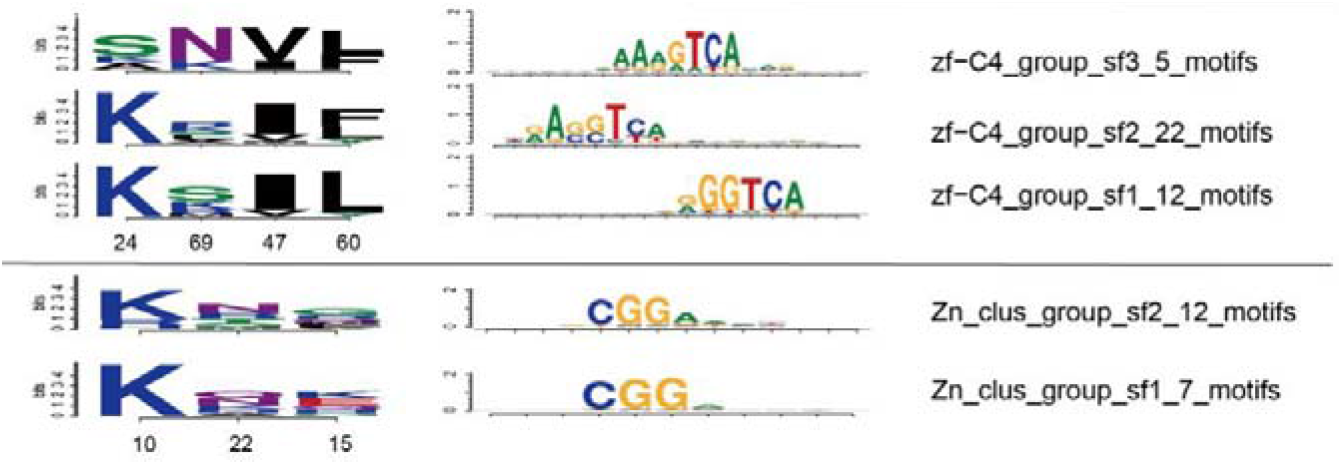
TF subgroup determining sites in subfamilies and corresponding DNA motifs.

**Figure S4.**
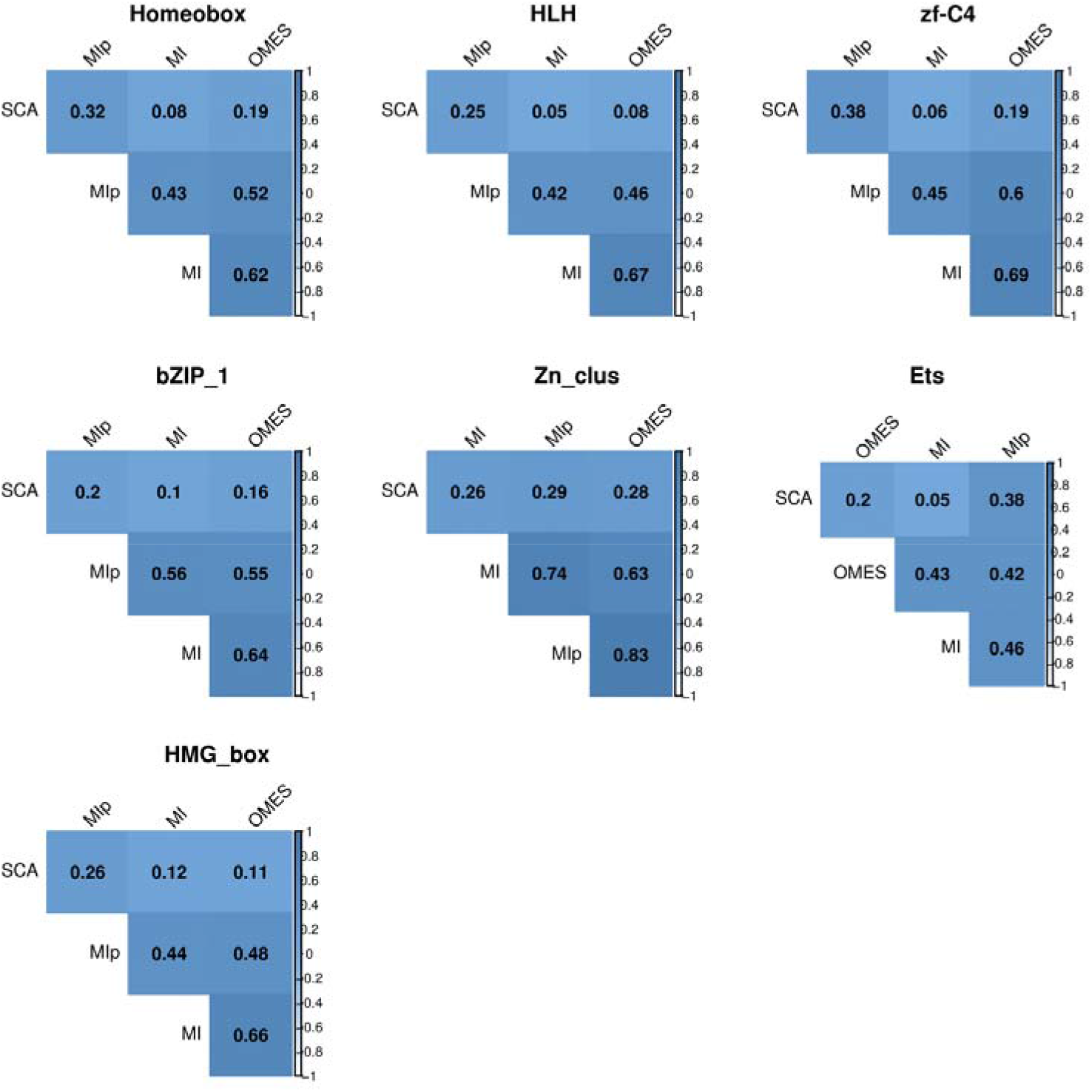
Jaccard-based correlation of four coevolution methods.

**Figure S5.**
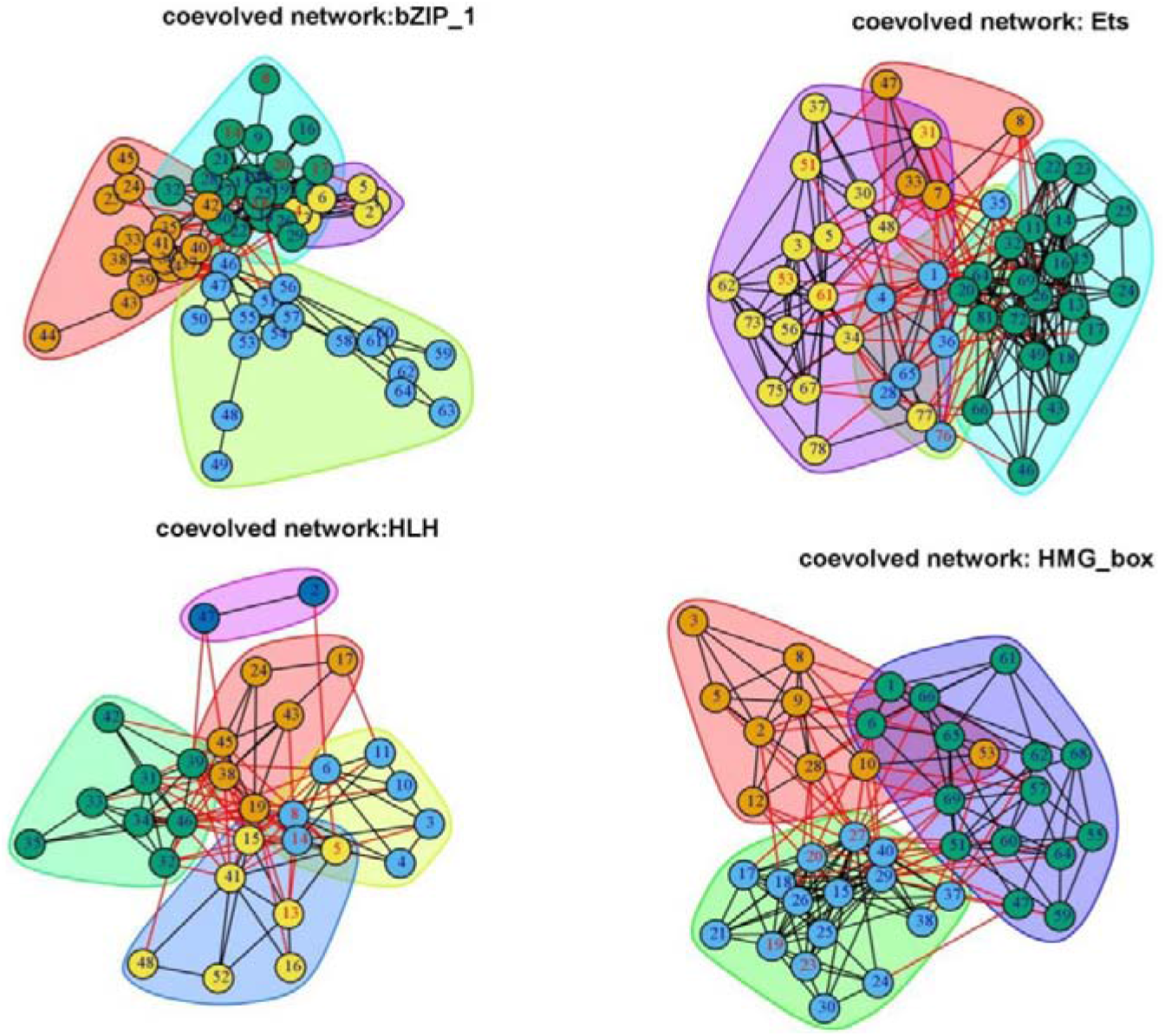
Network of coevolving residues of TF families, related to Figure 3E.

**Figure S6.**
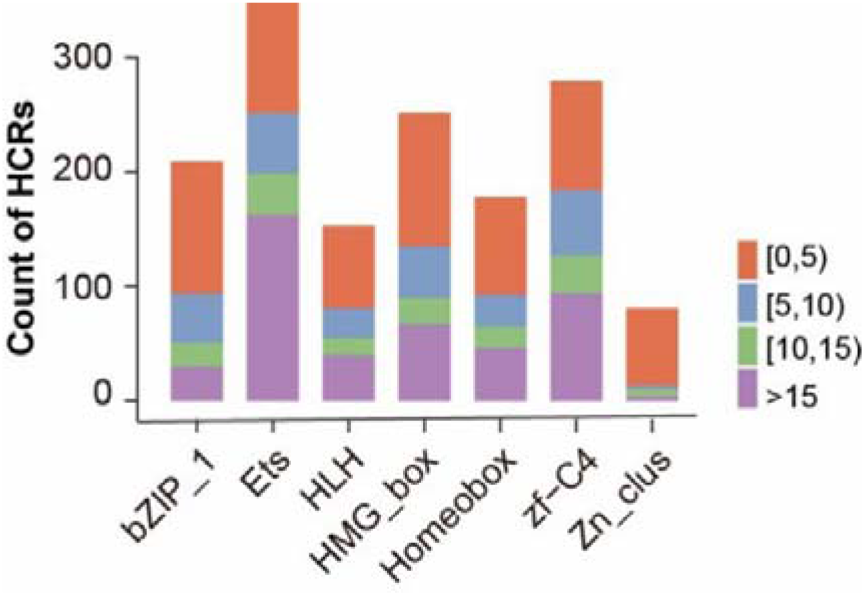
The coevolving residue distance within MSA profile.

**Figure S7.**
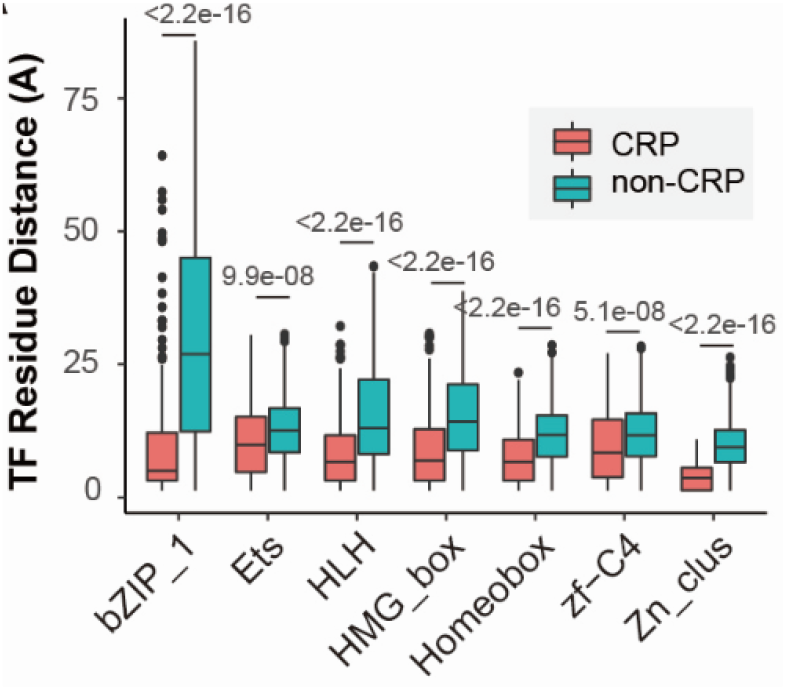
Comparing the distance between coevolving residue pairs (CRPs) and non-CRPs.

**Table S1.** Overview of included TF-DNA interaction assays.

| Source | Data type | PMID | # TFs | Description |
| --- | --- | --- | --- | --- |
| Berger_2006 | PBM | 16998473 | 4 | TFs from yeast, worm, mouse and human |
| Berger_2008 | PBM | 18585359 | 157 | Homeobox proteins in mouse |
| Badis_2008 | PBM | 19111667 | 110 | HLH, bZIP_1, zf, Fork_head, Homeobox in yeast |
| Scharer_2009 | PBM | 19147588 | 1 | HMG_box in human |
| Zhu_2009 | PBM | 19158363 | 29 | bZIP_1, HLH, zf, fork_head, HMG_box in yeast |
| Lesch_2009 | PBM | 19204119 | 1 | Homeobox in *C. elegans* |
| Badis_2009 | PBM | 19443739 | 101 | HLH, bZIP_1, zf, Ets, fork_head, HMG_box in mouse |
| Grove_2009 | PBM | 19632181 | 10 | HLH in *C. elegans* |
| Jolma_2013 | SELEX | 23332764 | 258 | HLH, bZIP_1, zf, Ets, fork_head, HMG_box, Homeobox in human |
| Weirauch MT_2014 | PBM | 25215497 | 932 | TFs in Arabidopsis, mouse, *C. elegans*, human and fly |
| FlyFactorSurvey | B1H | 21097781 | 297 | HLH, zf, bZIP_1, Ets, Homeobox, forkhead TFs in fly |
| Wei_2010 | PBM | 20517297 | 16 | Ets in mouse |

**Table S2.** Coevolving residue pairs. *Attached xls file (“S3.highly_coevolved_residues.xlsx”)*

**Table S3.** Residue blocks in TF domains.

| TF Family | coevolving residue positions |
| --- | --- |
| HLH | [1] "13_14_15_16"<br>[2] "1_2_3_4_5_6_7_8_9_10_11_12"<br>[3] "19_20_21"<br>[4] "29_30_31_32_33_34_35_36_37_38_39_40_41_42_43"<br>[5] "45_46_47_48" |
| zf-C4 | [1] "3_4_5_6_7_8_9_10_11_12_13_14_15_16_17_18_19_20_21_22_23"<br>[2] "30_31_32_33",<br>[3] "47_48_49_50_51_52_53_54"<br>[4] "54_55_56_57_58_59"<br>[5] "37_38_39" |
| bZIP_1 | [1] "1_2_3_4_5_6_7"<br>[2] "10_11_12"<br>[3] "23_24_25_26_27_28_29_30"<br>[4] "34_35_36_37_38_39_40_41_42_43_44_45_46_47"<br>[5] "53_54_55_56_57_58"<br>[6] "58_59_60_61_62_63_64"<br>[7] "17_18_19" |
| Zn_clus | [1] "14_15_16_17_18_19" "1_2_3_4_5_6_7_8_9"<br>[3] "24_25_26_27_28_29" "31_32_33_34_35_36_37_38_39_40" |
| Ets | [1] "30_31_32_33_34_35_36_37"<br>[2] "1_2_3_4_5_6_7_8_9_10_11_12_13_14_15_16_17_18_19_20"<br>[3] "23_24_25_26_27"<br>[4] "71_72_73_74_75_76_77_78_79_80_81"<br>[5] "61_62_63"<br>[6] "54_55_56"<br>[7] "65_66_67" |
| HMG_box | [1] "17_18_19_20_21_22_23_24_25_26_27_28_29_30_31_32_33"<br>[2] "51_52_53_54_55_56_57_58_59_60_61_62"<br>[3] "63_64_65_66_67_68_69"<br>[4] "1_2_3_4_5_6_7_8_9_10_11_12_13_14_15"<br>[5] "45_46_47" |

**Table S4.** Representative 3D structures of TF-DNA complex.

| TF Family | coevolving residue positions |
| --- | --- |
| HLH | 4H10, 1NLW, 1AN2, 1HLO, 1NKP, 4ATI, 4ATK, 1MDY, 2QL2, 1A0A, 1AM9, 2YPA, 2YPB, 1AN4 |
| zf-C4 | 1R4I, 1R0N, 1R0O, 2HAN, 1HCQ, 4AA6, 4HN5, 4HN6, 1GLU, 1LAT, 1R4O, 1R4R, 3FYL, 3G6P, 3G6Q, 3G6R, 3G6T, 3G6U, 3G8U, 3G8X, 3G97, 3G99, 3G9I, 3G9J, 3G9M, 3G9O, 3G9P, 3CBB, 4IQR, 4TNT, 1A6Y, 1GA5, 1HLZ, 1CIT, 2A66, 3DZU, 3DZY, 3E00, 1DSZ, 1BY4, 4CN2, 4CN3, 4CN5, 4CN7, 4NQA, 1YNW, 2NLL, 3M9E, 1KB2, 1KB4, 1KB6 |
| bZIP_1 | 1H8A, 1H89, 1H88, 1NWQ, 1JNM, 1GU5, 1GU4, 1GTW, 2E43, 2E42, 3A5T, 2WT7, 2WTY, 4AUW, 1HJC, 1GD2, 2DGC |
| Zn_clus | 1D66, 3COQ, 1HWT, 1QP9, 2HAP, 2ER8, 2ERE, 2ERG, 1PYI, 1ZME |
| Ets | 3JTG, 1DUX, 1BC7, 1BC8, 1HBX, 1K6O, 4IRI, 2NNY, 3MFK, 3RI4, 3WTS, 3WTT, 3WTU, 3WTV, 3WTW, 3WTX, 3WTY, 3WU1, 4L0Y, 4L0Z, 4L18, 4LG0, 1K78, 1K79, 1K7A, 1MDM, 4BQA, 4BNC, 4UUV, 4UNO, 4MHG, 3ZP5, 5E8I, 5JVT, 1AWC, 1YO5, 1PUE |
| HMG_box | 1QRV, 3NM9, 3F27, 4Y60, 3U2B, 4EUW, 4S2Q, 3TMM, 3TQ6, 4NNU, 4NOD |
| Homeobox | 3A01, 3LNQ, 9ANT, 4RDU, 1JGG, 1B8I, 2R5Y, 2R5Z, 4CYC, 4UUS, 4J19, 2HDD, 2HOS, 2HOT, 1DU0, 1HDD, 3HDD, 1IC8, 2H8R, 1PUF, 1B72, 1IG7, 1AKH, 1APL, 1MNM, 1YRN, 1K61, 1LE8, 4RBO, 3RKQ, 3CMY, 1AU7, 1CQT, 2XSD, 3L1P, 3D1N, 1FJL, 4S0H |

## Reference

1. Lambert, S.A., et al., The Human Transcription Factors. Cell, 2018. 172(4): p. 650–665.

2. Todeschini, A.L., A. Georges, and R.A. Veitia, Transcription factors: specific DNA binding and specific gene regulation. Trends Genet, 2014. 30(6): p. 211–9.

3. Slattery, M., et al., Absence of a simple code: how transcription factors read the genome. Trends Biochem Sci, 2014. 39(9): p. 381–99.

4. Ren, R., et al., Structural basis of specific DNA binding by the transcription factor ZBTB24. Nucleic Acids Res, 2019. 47(16): p. 8388–8398.

5. Rohs, R., et al., The role of DNA shape in protein-DNA recognition. Nature, 2009. 461(7268): p. 1248–53.

6. Gordan, R., et al., Genomic regions flanking E-box binding sites influence DNA binding specificity of bHLH transcription factors through DNA shape. Cell Rep, 2013. 3(4): p. 1093–104.

7. Kribelbauer, J.F., et al., Context-Dependent Gene Regulation by Homeodomain Transcription Factor Complexes Revealed by Shape-Readout Deficient Proteins. Mol Cell, 2020. 78(1): p. 152–167 e11.

8. Mukherjee, S., et al., Rapid analysis of the DNA-binding specificities of transcription factors with DNA microarrays. Nat Genet, 2004. 36(12): p. 1331–9.

9. Berger, M.F., et al., Variation in homeodomain DNA binding revealed by high-resolution analysis of sequence preferences. Cell, 2008. 133(7): p. 1266–76.

10. Meng, X., M.H. Brodsky, and S.A. Wolfe, A bacterial one-hybrid system for determining the DNA-binding specificity of transcription factors. Nat Biotechnol, 2005. 23(8): p. 988–94.

11. Jolma, A., et al., Multiplexed massively parallel SELEX for characterization of human transcription factor binding specificities. Genome Res, 2010. 20(6): p. 861–73.

12. Slattery, M., et al., Cofactor binding evokes latent differences in DNA binding specificity between Hox proteins. Cell, 2011. 147(6): p. 1270–82.

13. Weirauch, M.T., et al., Determination and inference of eukaryotic transcription factor sequence specificity. Cell, 2014. 158(6): p. 1431–1443.

14. Badis, G., et al., Diversity and complexity in DNA recognition by transcription factors. Science, 2009. 324(5935): p. 1720–3.

15. Jolma, A., et al., DNA-binding specificities of human transcription factors. Cell, 2013. 152(1–2): p. 327–39.

16. Chakrabarti S. and A.R. Panchenko, Coevolution in defining the functional specificity. Proteins, 2009. 75(1): p. 231–40.

17. Chakrabarti S. and A.R. Panchenko, Structural and functional roles of coevolved sites in proteins. PLoS One, 2010. 5(1): p. e8591.

18. Clark, G.W., et al., Using coevolution to predict protein-protein interactions. Methods Mol Biol, 2011. 781: p. 237–56.

19. Wang, Y., et al., Coevolution-based prediction of protein-protein interactions in polyketide biosynthetic assembly lines. Bioinformatics, 2020.

20. Sutto, L., et al., From residue coevolution to protein conformational ensembles and functional dynamics. Proc Natl Acad Sci U S A, 2015. 112(44): p. 13567–72.

21. Toth-Petroczy, A., et al., Structured States of Disordered Proteins from Genomic Sequences. Cell, 2016. 167(1): p. 158–170 e12.

22. Reimer, J.M., et al., Structures of a dimodular nonribosomal peptide synthetase reveal conformational flexibility. Science, 2019. 366(6466).

23. Sfriso, P., et al., Residues Coevolution Guides the Systematic Identification of Alternative Functional Conformations in Proteins. Structure, 2016. 24(1): p. 116–126.

24. Ribeiro, A.J.M., et al., A global analysis of function and conservation of catalytic residues in enzymes. J Biol Chem, 2020. 295(2): p. 314–324.

25. Petrovic, D., et al., Conformational dynamics and enzyme evolution. J R Soc Interface, 2018. 15(144).

26. Chan, T.M., et al., Subtypes of associated protein-DNA (Transcription Factor-Transcription Factor Binding Site) patterns. Nucleic Acids Res, 2012. 40(19): p. 9392–403.

27. Yang, S., et al., Correlated evolution of transcription factors and their binding sites. Bioinformatics, 2011. 27(21): p. 2972–8.

28. Laforet, M., et al., Modifying a covarying protein-DNA interaction changes substrate preference of a site-specific endonuclease. Nucleic Acids Res, 2019. 47(20): p. 10830–10841.

29. Hertz, G.Z., G.W. Hartzell, 3rd, and G.D. Stormo, Identification of consensus patterns in unaligned DNA sequences known to be functionally related. Comput Appl Biosci, 1990. 6(2): p. 81–92.

30. Ou, J., et al., motifStack for the analysis of transcription factor binding site evolution. Nat Methods, 2018. 15(1): p. 8–9.

31. Mistry, J., et al., Pfam: The protein families database in 2021. Nucleic Acids Res, 2020.

32. Mercier, E., et al., An integrated pipeline for the genome-wide analysis of transcription factor binding sites from ChIP-Seq. PLoS One, 2011. 6(2): p. e16432.

33. Edgar, R.C., MUSCLE: multiple sequence alignment with high accuracy and high throughput. Nucleic Acids Res, 2004. 32(5): p. 1792–7.

34. Chakrabarti, S., S.H. Bryant, and A.R. Panchenko, Functional specificity lies within the properties and evolutionary changes of amino acids. J Mol Biol, 2007. 373(3): p. 801–10.

35. Gloor, G.B., et al., Mutual information in protein multiple sequence alignments reveals two classes of coevolving positions. Biochemistry, 2005. 44(19): p. 7156–65.

36. Dunn, S.D., L.M. Wahl, and G.B. Gloor, Mutual information without the influence of phylogeny or entropy dramatically improves residue contact prediction. Bioinformatics, 2008. 24(3): p. 333–40.

37. Lockless S.W. and R. Ranganathan, Evolutionarily conserved pathways of energetic connectivity in protein families. Science, 1999. 286(5438): p. 295–9.

38. Fodor A.A. and R.W. Aldrich, Influence of conservation on calculations of amino acid covariance in multiple sequence alignments. Proteins, 2004. 56(2): p. 211–21.

39. Bakan, A., et al., Evol and ProDy for bridging protein sequence evolution and structural dynamics. Bioinformatics, 2014. 30(18): p. 2681–3.

40. Clauset, A., M.E. Newman, and C. Moore, Finding community structure in very large networks. Phys Rev E Stat Nonlin Soft Matter Phys, 2004. 70(6 Pt 2): p. 066111.

41. Burley, S.K., et al., RCSB Protein Data Bank: biological macromolecular structures enabling research and education in fundamental biology, biomedicine, biotechnology and energy. Nucleic Acids Res, 2019. 47(D1): p. D464–D474.

42. Schymkowitz, J., et al., The FoldX web server: an online force field. Nucleic Acids Res, 2005. 33(Web Server issue): p. W382–8.

43. Hu, H., et al., AnimalTFDB 3.0: a comprehensive resource for annotation and prediction of animal transcription factors. Nucleic Acids Res, 2019. 47(D1): p. D33–D38.

44. Lambert, S.A., et al., Similarity regression predicts evolution of transcription factor sequence specificity. Nat Genet, 2019. 51(6): p. 981–989.

45. Siggers, T., et al., Diversification of transcription factor paralogs via noncanonical modularity in C2H2 zinc finger DNA binding. Mol Cell, 2014. 55(4): p. 640–8.

46. Noyes, M.B., et al., Analysis of homeodomain specificities allows the family-wide prediction of preferred recognition sites. Cell, 2008. 133(7): p. 1277–89.

47. Treisman, J., et al., A single amino acid can determine the DNA binding specificity of homeodomain proteins. Cell, 1989. 59(3): p. 553–62.

48. Grove, C.A., et al., A multiparameter network reveals extensive divergence between C. elegans bHLH transcription factors. Cell, 2009. 138(2): p. 314–27.

49. De Masi, F., et al., Using a structural and logics systems approach to infer bHLH-DNA binding specificity determinants. Nucleic Acids Res, 2011. 39(11): p. 4553–63.

50. Wei, G.H., et al., Genome-wide analysis of ETS-family DNA-binding in vitro and in vivo. EMBO J, 2010. 29(13): p. 2147–60.

51. Malarkey C.S. and M.E. Churchill, The high mobility group box: the ultimate utility player of a cell. Trends Biochem Sci, 2012. 37(12): p. 553–62.

52. Fujii, Y., et al., Structural basis for the diversity of DNA recognition by bZIP transcription factors. Nat Struct Biol, 2000. 7(10): p. 889–93.

53. Colell, E.A., et al., MISTIC2: comprehensive server to study coevolution in protein families. Nucleic Acids Res, 2018. 46(W1): p. W323–W328.

54. Mao, W., et al., Comparative study of the effectiveness and limitations of current methods for detecting sequence coevolution. Bioinformatics, 2015. 31(12): p. 1929–37.

55. Dib L. and A. Carbone, Protein fragments: functional and structural roles of their coevolution networks. PLoS One, 2012. 7(11): p. e48124.

56. Anishchenko, I., et al., Origins of coevolution between residues distant in protein 3D structures. Proc Natl Acad Sci U S A, 2017. 114(34): p. 9122–9127.

57. Phelan M.L. and M.S. Featherstone, Distinct HOX N-terminal arm residues are responsible for specificity of DNA recognition by HOX monomers and HOX.PBX heterodimers. J Biol Chem, 1997. 272(13): p. 8635–43.

